# Aging Alters Hair Cell Physiological Properties in Mice with Late-Onset Age-Related Hearing Loss

**DOI:** 10.1101/2025.10.24.684339

**Authors:** Piece Yen, Adam J. Carlton, Andrew O’Connor, Stuart L. Johnson, Jing-Yi Jeng

**Affiliations:** School of Biomedical Science, University of Sheffield, Sheffield, S10 2TN, UK; Neuroscience Institute, University of Sheffield, Sheffield, S10 2TN, UK; Wellcome Sanger Institute, Hinxton, Cambridge, CB10 1SA, UK

**Keywords:** Hair cell physiology, age-related hearing loss, cochlear aging

## Abstract

Cochlear hair cells convert sound stimuli into electrical signals. Age-related hearing loss (ARHL) is characterized by the progressive reduction of these hair cells and their synapses, but the underlying mechanisms remain poorly understood.

In this study, we tested the hypothesis that hair cell function declines with age in CBA/CaJ mice, a strain known for its very slow ARHL progression. We found that hair cell size began to decrease from 10 months of age, before hearing loss was detectable. This structural change was associated with a significant decrease in the total current response and a distinct reduction in the expression of large conductance calcium-activated potassium (BK) channel currents (*I*_K,f_). However, the magnitude of the mechanoelectrical transducer (MET) currents at the hair cell stereociliary bundles remained unaffected. This observation challenges the common conception that stereocilia are the most fragile component of sensory cells in aging.

Our results provide clear evidence of an early decline in hair cell function with age, which precedes measurable hearing loss, suggesting that these cellular deficits define a distinct pathological route to ARHL in the CBA/CaJ mouse model.

## Introduction

ARHL, also known as presbycusis, is one of the most common age-related disorders affecting over one third of adults over 60 years old (Cieza et al., 2021; *World Report on Hearing*, 2021). Despite its high prevalence, the underlying mechanisms of ARHL are still unclear due to its complexity. The progression of ARHL is highly variable in humans, starting between 30 and 70 years of age (Yamasoba et al., 2013). It involves various pathologies, which can differ among individuals, including the degeneration of spiral ganglion neurons (SGNs) and the sensory hair cells within the cochlea (Gacek & Schuknecht, 1969; P. Z. Wu et al., 2018). Furthermore, ARHL is linked to risk factors such as genetic predispositions, environmental insults, and lifestyle choices (Gioacchini et al., 2023; Tang et al., 2023; P.-Z. Wu et al., 2021). To study these mechanisms, various mouse models represent different patterns of age-related hearing loss. For example, C57BL/6N mice model early-onset ARHL, while C3H and CBA mice model late-onset ARHL. Although the factors contributing to the variability within and among these models are still unclear, they generally share similar underlying pathologies.

A common pathology of ARHL is the loss of the two types of sensory cells in the mammalian cochlea: inner hair cells (IHCs) and outer hair cells (OHCs). OHCs provide acoustic amplification, while IHCs are the primary sound-transducing sensors. Both cell types are activated when sound-induced deflection of their stereociliary bundles opens the mechanically-sensitive mechanoelectrical transducer (MET) channels located at the tips of the shorter stereociliary rows (Assad et al., 1991; Beurg et al., 2009). As MET channels open, an influx of cations depolarizes the cell, generating a receptor potential that triggers the release of synaptic vesicles onto SGN terminals. This receptor potential is a graded and sustained response, modulated by K^+^ currents that are crucial for regulating the resting membrane potential and preventing action potentials. Following the onset of hearing, hair cells express characteristic K^+^ currents: *I*_K,n_ and *I*_K,f_. *I*_K,n_ is a negatively-activating rectifying current carried out by KCNQ family K^+^ channels, and *I*_K,f_ is a fast K^+^ current mediated by the activation of large-conductance Ca^2+^-activated K^+^ channels (BK channels) (Kros et al., 1998; Marcotti et al., 2004b; Marcotti & Kros, 1999; Oliver et al., 2003). Across all cochlear regions, OHCs predominantly express *I*_K,n_, while IHCs predominately express *I*_K,f_ (Rohmann et al., 2015; Wersinger et al., 2010; Johnson, 2015). In an early-onset ARHL model, C57BL/6N mice, hair cells show reduced *I*_K,f_, MET currents and cell size (Jeng, Johnson, et al., 2020; Jeng, Carlton, et al., 2021), suggesting that hair cells degenerate functionally with age. However, it is unclear if this degeneration occurs across all ARHL mouse models.

In contrast to C57BL/6N mice, CBA mice develop a much later-onset ARHL, with significant hearing loss occurring after one year of age, making them a standard late-onset ARHL model (Ohlemiller et al., 2010; Sha et al., 2008; Spongr et al., 1997). Crucially, microscopic analysis has well-established that IHCs in CBA mice begin to lose their synaptic ribbons before hearing threshold changes are evident, a phenomenon known as hidden hearing loss (Sergeyenko et al., 2013; Liberman & Kujawa, 2017; Parthasarathy & Kujawa, 2018; Dörje et al., 2024). This suggests a distinct pathological mechanism within IHCs that triggers synapse loss and contributes to later hearing decline.

To test the hypothesis that hair cells undergo functional degeneration preceding the onset of ARHL, we investigated the physiological properties of IHCs and OHCs from CBA/CaJ mice using patch-clamp electrophysiology and imaging. We found that hair cells are smaller with age and that IHCs lose their *I*_K,f_ started much earlier before the onset of hearing threshold shift. Unlike previous observations in C57BL/6N mice, we found no evidence of stereociliary degeneration via electron microscopy, nor did the size of their MET currents decrease with age. As was previously reported in C57BL/6N mice (Zachary & Fuchs, 2015; J. Jeng et al., 2020), the IHCs of CBA/CaJ mice were also innervated by efferent neurons, although this process began much later, following the changes in the IHC current profile. These findings collectively underscore that physiological degeneration is a key, early factor involved in the pathogenesis of ARHL.

## Results

### CBA/CaJ mice show hidden hearing loss with age

We first determined the hearing thresholds in aging CBA/CaJ mice by measuring auditory brainstem responses (ABRs). As expected, hearing thresholds measured from ABRs show only mild hearing loss with age (Figure 1A–C). In aging CBA/CaJ mice, the ABR thresholds were elevated significantly when stimulated with a click or a pure tone at 36 kHz at 21–23 months (Figure 1A, P = 0.031 and 0.028, respectively).

**Figure 1.**
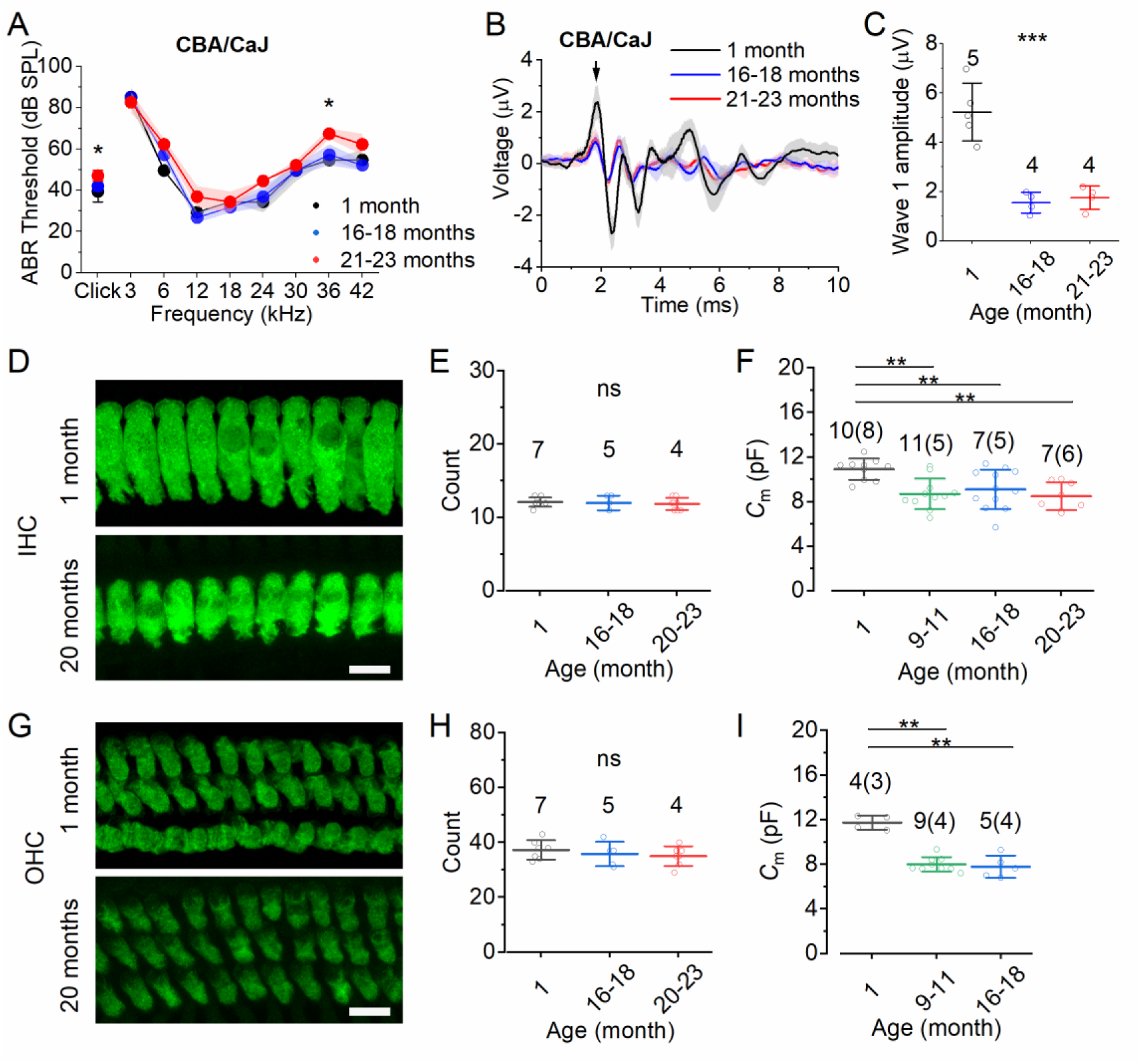
Hair cells become smaller with age in hidden hearing loss mice. *A–C*, ABR recordings for click and pure-tone stimulation of 3–42 kHz recorded from CBA/CaJ mice at 1, 16–18 and 21–23 months. ABR thresholds from CBA/CaJ mice are shown as median ± median absolute deviation in A, *P < 0.05, Kruskal-Wallis ANOVA. Average waveform responses recorded at 12 kHz and stimulated at 70 dB from CBA/CaJ mice are shown as mean ± SD (B). Average amplitude of wave 1 (arrows in B and E) are plotted as mean ± SD (C). Single-animal recordings are plotted as open symbols. The number of CBA/CaJ animals in A–C is shown in C. ***P < 0.001, **P = 0.011, one-way ANOVA. *D*–*I*, Quantification of the number of cells and cell membrane size of IHCs (D–F) and OHCs (G–I) in ageing CBA/CaJ mice. Maximum intensity projections of confocal z-stacks of inner hair cells (IHCs) (D) and outer hair cells (OHCs) (G) from CBA/CaJ mice at 1 month and 20 months. The cells were labelled with antibodies against a hair cell marker MYO7A (green). The number of cells were counted across a 100 µm region from the apical coil of mice at 1, 16– 18 and 20–23 months, and the values from each mouse were plotted behind the mean ± SD values for IHCs (E) and OHCs (H). For cell membrane size, average membrane capacitance (*C*m) values were recorded from apical IHCs (F) and OHCs (I), and recordings from each cell are shown behind the mean ± SD values. The number of cells (animals) are indicated above each data set. ** P < 0.01, one-way ANOVA followed by Tukey’s post-hoc test, ns: not significant Scale bar: 10 µm.

To further assess the output from the cochlea, we analysed ABR wave 1 generated by the auditory nerve that exits the cochlea. We selected wave 1 recorded at 75 dB at 12 kHz where there was no significant hearing threshold shift in CBA/CaJ and C3H/HeJ mice (arrows, Figure 1B and E). Despite no threshold shift, both mouse strains showed decreased wave 1 peak-to-peak amplitude with age, suggesting a reduction in the output from the auditory nerve (CBA/CaJ, P < 0.001; C3H/HeJ, P = 0.011, Figure 1C and F). Overall, CBA/CaJ mice present a mild hearing loss with age, showing hidden hearing loss at 12 kHz at old ages.

### Hair cells become smaller with age in CBA/CaJ mice

The reduction in the wave 1 amplitude in CBA/CaJ mice suggests a problem with the peripheral auditory system, including the auditory neurons and the hair cells that transduce the signal to the auditory nerves. To confirm that there is no hair cell loss at this age, we quantified the number of hair cells with an antibody against myosin 7a (MYO7A), which is used as a hair cell marker (IHC: Figure 1D, and OHC: Figure 1G). Indeed, there was no significant hair cell loss across a 100 µm region at the apical turn of the cochlea before 23 months of age (IHC: Figure 1E, and OHC: Figure 1H). This result is similar to that reported by other groups previously in CBA/CaJ mice (Dörje et al., 2024; Sergeyenko et al., 2013). However, we noticed that IHCs appeared to be much smaller at old ages from the MYO7A staining (Figure 1A). To accurately quantify the size of hair cells, we performed patch-clamp recordings to measure the total cell membrane capacitance (*C*_m_), which is proportional to the cell surface area (Solsona et al., 1998). We found that not only IHCs but also OHCs showed a significant decrease in *C*_m_ from ages as early as 9-month-compared to 1-month-old at the apical coil of the cochlea (9–12 kHz) (IHC: 9–11 months P = 0.005, 16–18 months P = 0.023, 20–23 months P = 0.004, Figure 1C; OHC: 9–11 months P < 0.001, 16–18 months P < 0.001, Figure 1F). Taken together, these data indicate that hair cell loss is not present in the apex of CBA/CaJ animals. However, aging hair cells become smaller with age. One theory is that smaller cell size is usually associated with a reduced expression of ion channels to maintain current density and the function of the cell (Gorur-Shandilya et al., 2020).

### The rapidly-activating K+ current in IHCs becomes smaller in aging CBA/CaJ mice

To determine whether hair cell function changes in the progression of hidden hearing loss, we performed whole-cell patch-clamp to analyse physiological changes from aging hair cells at the apical coil of the cochlea (9–12 kHz). At this frequency range, we were able to reliably record single-cell electrophysiology as a detailed readout of hair cell function.

Current responses from voltage-gated K^+^ channels were elicited using a series of hyperpolarizing and depolarizing voltage steps from a holding potential of -84 mV (Figure 2). For OHCs, we measured the size of the total steady-state K^+^ current *I*_K_ at 0 mV and the deactivating current *I*_K,n_ at 1.5 ms from the current onset at -124 mV (Figure 2A and B). With age, CBA/CaJ mice expressed less total current in OHCs (1 vs. 16–18 month, P = 0.014, Figure 2C). To investigate whether the reduction in current is related to smaller cell size, we calculated cell current density by normalising currents to their respective *C*_m_. The resulting current density was not significantly different with age, indicating that the reduction in *I*_K_ is related to cell shrinkage (P = 0.452, Figure 2D). In mature mouse OHCs, more than half of *I*_K_ is composed of a negatively-activating current *I*_K,n_, which is important for maintaining a negative resting membrane potential (Kharkovets et al., 2006; Marcotti & Kros, 1999). Compared to *I*_K_, the size of *I*_K,n_ was well-regulated with age and there was no significant difference in the current density (P = 0.629 for *I*_K,n_ and P = 0.400 for *I*_K,n_/*C*_m_, Figure 2E and F).

**Figure 2.**
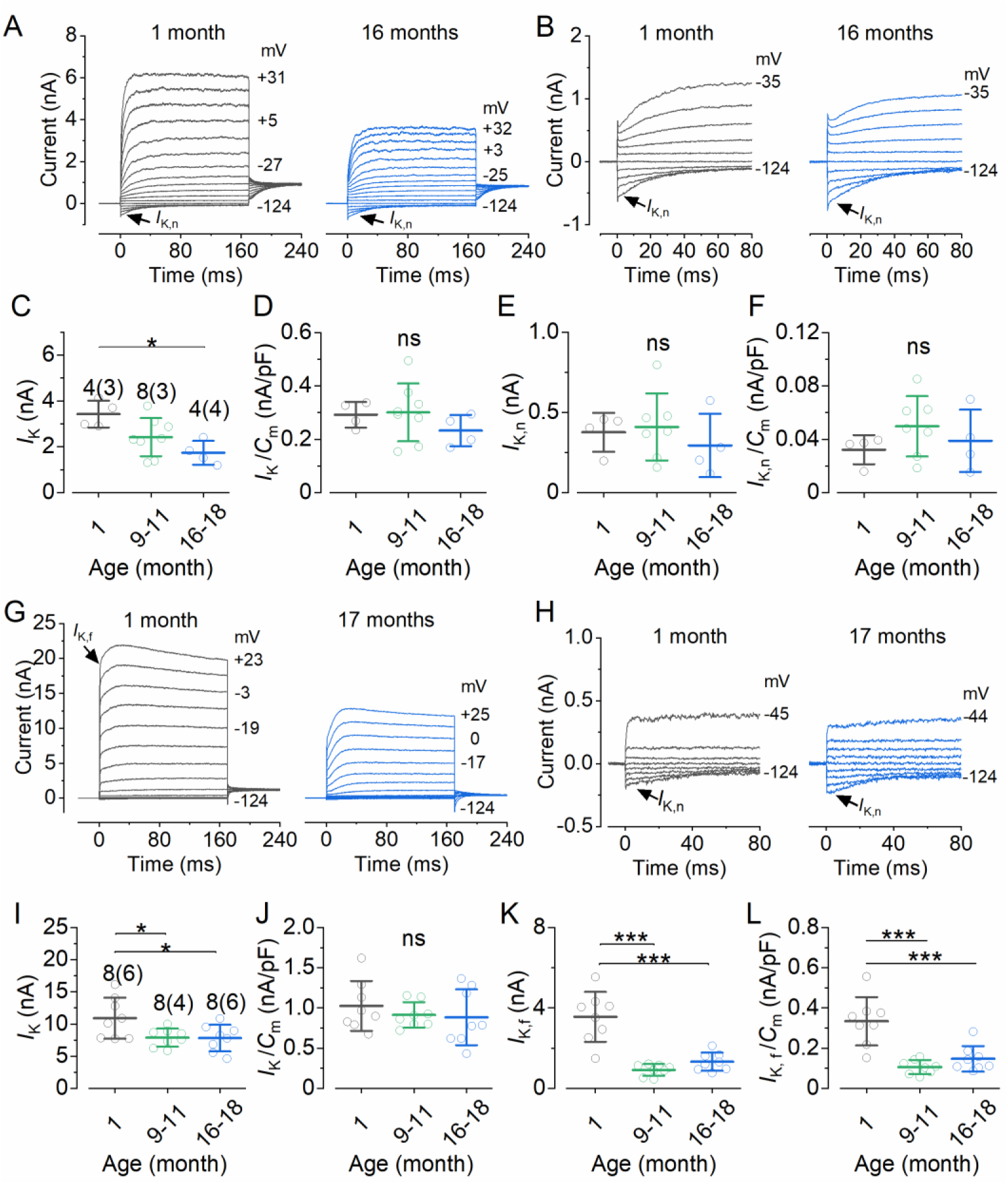
Hair cell current responses decrease with age. Potassium current recorded from ageing OHCs (A–F) and IHCs (G–L) in CBA/CaJ mice. Cells were held at -84 mV and currents were recorded using 10 mV depolarising voltage steps from -124 mV. Various test potentials are shown by some of the traces. *A* and *B*, examples of K+ currents recorded from OHCs at 1 and 16 months. The presence of *I*K,n is indicated at the beginning of the trace. *C*, size of peak K+ current from OHCs measured at a membrane potential of 0 mV. *D,* current density measured by normalising peak *I*K (C) to the corresponding cell membrane capacitance (*C*m). *E*, size of *I*K,n measured as the deactivating currents at -124 mV. *F*, current density measured by normalising *I*K,n (E) to the corresponding *C*m. *G* and *H*, examples of K+ currents recorded from IHCs at 1 and 17 months. The presence of *I*K,f and *I*K,n is indicated at the beginning of the trace. *I*, size of peak K+ current from IHCs measured at a membrane potential of 0 mV. *J,* current density measured by normalising peak *I*K (I) to the corresponding cell membrane capacitance (*C*m). *K*, size of *I*K,f measured as the deactivating currents at -124 mV. *L*, current density measured by normalising *I*K,f (K) to the corresponding *C*m. Single-cell recordings are plotted as open symbols. The number of cells (animals) in C–F and I–L is shown above the data points in C and I, separately. * P < 0.05, *** P < 0.001, one-way ANOVA followed by Tukey’s post-hoc test.

For mature mouse IHCs, the total current appears to reduce with age (Figure 2G). To quantify this, the size of steady-state *I*_K_ at 0 mV was measured. The results show that *I*_K_ was significantly reduced with age without changing their current density (*I*_K_ at 1 month vs. 9–11 months P = 0.044; vs. 16–18 months P = 0.038, and P = 0.583 for *I*_K_/*C*_m_, Figure 2I). In mature IHCs, *I*_K_ is composed of *I*_K,s_, *I*_K,n_ and *I*_K,f_. To further investigate whether *I*_K,n_ and *I*_K,f_ contribute the decline of *I*_K_, we measured separately *I*_K,n_ at -124 mV and *I*_K,f_ at -25 mV, at 1.5 ms from the current onset at (Jeng, Carlton, et al., 2021b; Jeng et al., 2022). Like OHCs, there was no significant change in *I*_K,n_ with age (1 month: 0.109 ± 0.027 nA, 9–11 months: 0.112 ± 0.051 nA, 16–18 months: 0.100 ± 0.037 nA, P = 0.811, one-way ANOVA). On the other hand, the size of *I*_K,f_ and its current density significantly decreased with age (aging vs. 1 month: P < 0.001 for *I*_K,f_ and *I*_K,f_/*C*_m_ at 9–11 and 16–18 months, Figure 2K and L). As *I*_K,f_ is carried by BK channel, this suggests there was less active BK channels in aging IHCs. Overall, these data indicate that aging hair cells changed their total K^+^ current profiles to compensate the decreasing cell size but IHCs were not able to maintain their rapid-activating K^+^ responses.

### The expression of BK channels in IHCs is reduced in aging CBA/CaJ mice

Considering the reduction in *I*_K,f_ with age, we further investigated the expression of BK channels that carry *I*_K,f_ in IHCs using immunolabeling. The experiment was also done in C57BL/6N mice as they also express reduced *I*_K,f_ with age compared to C3H mice (Beurg et al., 2018; Jeng, Carlton, et al., 2021b). BK channels were present at the neck region of the IHCs from aging CBA/CaJ, C57BL/6N and C3H/HeJ mice (Figure 3A and B, Lingle et al., 2019). By quantifying all puncta per cell using the Arivis software, we found the total volume of anti-BK signals for each cell was significantly decreased in aging animals (CBA/CaJ at 1 month vs. CBA/CaJ at 20–23 months: P < 0.001; vs. C3H at 20–23 months: P < 0.001; vs. 6N at 16–18 months: P < 0.001, Figure 3C).

**Figure 3.**
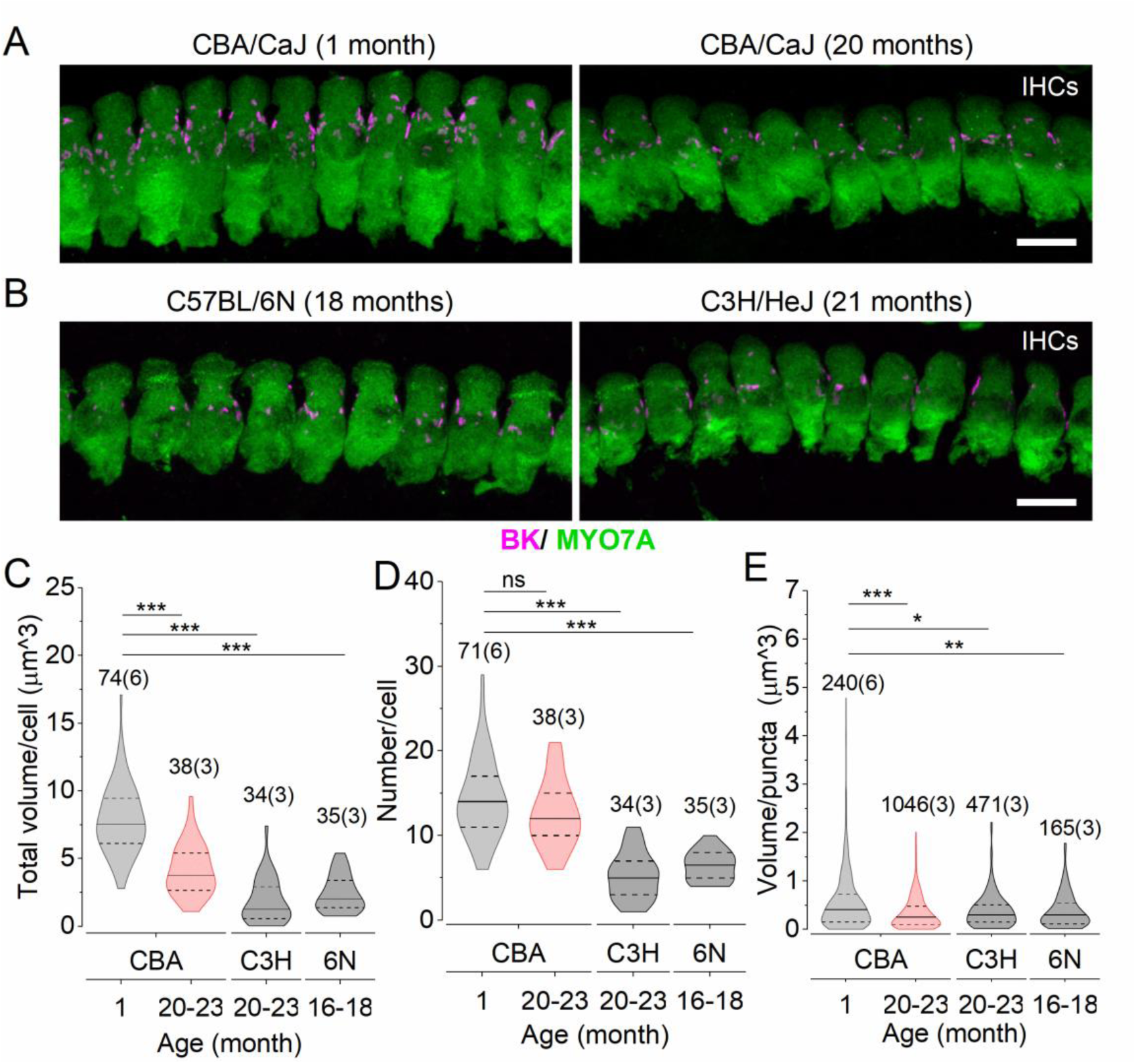
Expression of BK channels was reduced from ageing IHCs. Cochlea from the apical coil (9–12 kHz) of CBA/CaJ (CBA), C3H/HeJ (C3H) and C57BL/6N (6N) mice were labelled with antibodies against a hair cell marker MYO7A (green) and BK channels (red). *A* and *B*, maximum intensity projections of confocal z-stacks of IHCs from a CBA/CaJ mouse at 1 month and 20 months, a C57BL/6N mouse at 18 months, and a C3H/HeJ mouse at 21 months. *C*, total volume of BK puncta per IHC. Number of cells is shown above the data. *D*, total number of BK puncta per IHC. Number of cells is shown above the data. *E*, volume of each single BK puncta. Number of puncta is shown above the data. Number of animals is shown above the data in brackets in C-E. Scale bar: 10 µm. * P < 0.05, *** P < 0.001, ns: not significant, Kruskal-Wallis followed by Dunn’s post-hoc test.

To test whether the decrease of BK expression is caused by a reduced number of puncta, we analysed each puncta separately. Interestingly, we found no significant change in the number of puncta with age in CBA/CaJ but significantly less in C3H/HeJ and C57BL/6N mice (CBA/CaJ at 1 month vs. CBA/CaJ at 20–23 months: P > 0.999; vs. C3H/HeJ at 20–23 months: P < 0.001, vs. C57BL/6N at 16–18 months: P < 0.001, Figure 3D). This suggests that aging CBA/CaJ IHCs have a different aging process compared to C3H/HeJ and C57BL/6N mice. However, average BK volume for a single punctum was reduced in all aging mice, (CBA/CaJ at 1 month vs. CBA/CaJ at 20–23 months: P < 0.001, vs. C3H/HeJ at 20–23 months: P = 0.014, vs. C57BL/6N at 16–18 months: P = 0.001, Figure 3E). In CBA/CaJ IHCs, the decreasing *I*_K,f_ with age is associated with less BK channel expression but not BK channel localisation on the membrane.

### The mechanoelectrical transducer current is maintained in aging CBA/CaJ IHCs

As one can argue that the decrease in wave 1 amplitude is due to decreased mechanoelectrical transduction, we next investigated MET apparatus in aging CBA/CaJ mice. Using scanning electron microscopy, the stereociliary bundles in IHCs and OHCs had uniformly stair-case structure, and we did not see obvious morphological defect (Figure 4A–F). To test if mechanoelectrical transduction is unaffected with age, we were able to perform challenging MET recordings from aging IHCs (Jeng, Carlton, et al., 2021b). Example recordings from 1-, 17- and 22-month-old mice show that MET currents were elicited in aging IHCs by using a fluid-jet to deliver a sinusoidal displacement stimulus (Figure 4G–I).

**Figure 4.**
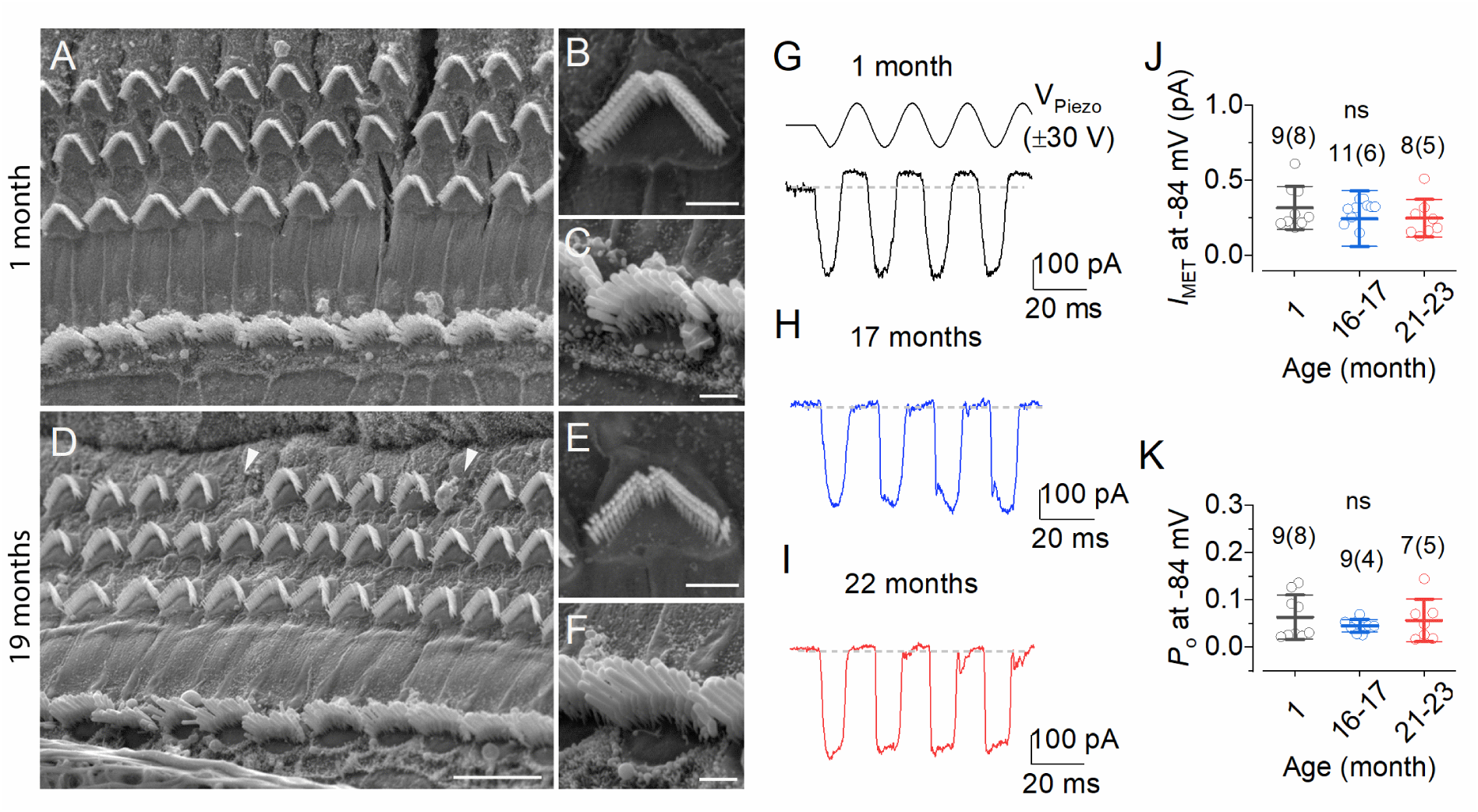
Mechanoelectrical transducer (MET) apparatus in ageing hair cells. *A-F*, scanning electron microscopy (SEM) showing the gross morphology of apical-coil hair cells from CBA/CaJ mice at 1 (A-C) or 19 (D-F) months. Some OHCs were missing at 19 months (arrowheads) but no obvious morphological changes were found in stereociliary bundles from outer hair cells (B and E) and inner hair cells (C and F). *G–I*, examples of saturating MET currents recorded from IHCs of CBA/CaJ mice at 1 (G), 17 (H) and 22 (I) months. MET currents were triggered by applying sinusoidal stimuli of 50 Hz to hair bundles from a holding potential of -84 mV. *J*, maximal size (J) and resting open probability (K) measured from MET current recordings in CBA/CaJ mice with age. Number of cells is shown above the data. Single-cell recordings are plotted as open symbols. ns: not significant, one-way ANOVA.

We next measured the maximum size and the open probability of the MET current. The maximum size of the MET current at a holding potential of -84 mV was measured from the current difference between the peak current during inhibitory and excitatory bundle displacement. The results show that the maximum MET current size was similar with age (P = 0.549, Figure 4J). The resting open probability (*P*_o_) of the MET current was then measured at a holding potential of -84 mV as the proportion of MET current active at rest. also similar with age (P = 0.591, Figure 4K). These recordings suggest that the MET apparatus in CBA/CaJ mice does not deteriorate with age, and as such does not contribute to ARHL in these animals.

Since the receptor potential is modulated by the opening of MET channels and the basolateral membrane currents, age-related decrease in *I*_K_ with preserved MET current in IHCs is likely to disrupt the resting membrane potential in CBA/CaJ mice. We thus measured resting membrane potential in IHCs with age, as well as investigating their voltage responses by injected currents from the resting membrane potential. This elicits fast and graded voltage responses in hair cells (Figure 5A–F). The resting membrane potential (*V*_m_) was measured at 0 current injection, and the results show that the resting *V*_m_ increased with age in IHCs but not OHCs (IHC 1 vs. 16–18 months: P = 0.007, OHC: P = 0.191, Figure 5G). It was noticeable that the resting *V*_m_ in aging OHCs was more variable compared to IHCs (variance at 16–18 months IHC: 4.811; OHC: 18.878). We further measured the voltage responses elicited with 0.9 nA current injection. Similar to the 0 current injection results, the 0.9 nA current injection triggered a higher *V*_m_ in aging IHCs but not OHCs (IHC 1 vs. 9–11 months: P = 0.008, IHC 1 vs. 16–18 months: P = 0.006, OHC: P = 0.484, Figure 5H). These data together provide evidence that aging CBA/CaJ IHCs maintained their MET current but had more depolarised resting membrane potential *ex vivo*, potentially due to a decreased overall *I*_K_.

**Figure 5.**
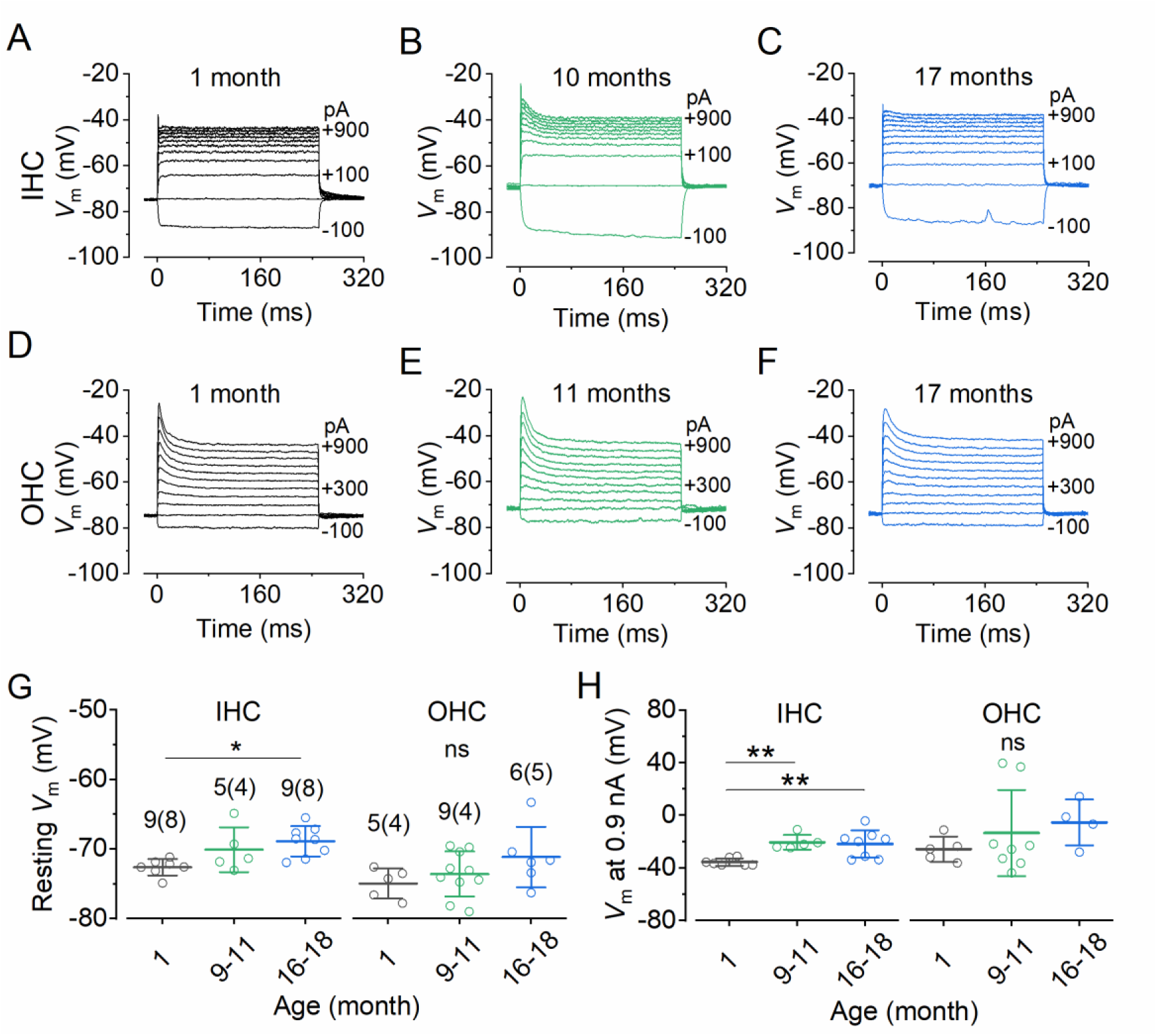
Ageing IHCs become more depolarised at rest. *A-F*, examples of voltage responses elicited by applying a series of 0.1 nA current injections to inner hair cells (IHCs) (A-C) and outer hair cells (OHCs) (D-F) from CBA/CaJ mice at 1, 10 and 17 months. *G*, resting membrane potential (*V*m) recorded from 1-, 9–11 and 16–18-month-old CBA/CaJ mice. *H,* voltage responses in hair cells elicited with 0.9 nA current injection in CBA/CaJ mice at 1, 9–11 and 16–18 months. Single-cell recordings are plotted as open symbols. The number of cells and animals in G and H is shown above the data points in G. The number of animals is in brackets. **P < 0.01, * P < 0.05, one-way ANOVA followed by Tukey’s post-hoc test, ns: not significant.

### Efferent fibres re-innervated aging IHCs

It was proposed that reduced mechanoelectrical transduction from hair cells leads to efferent fibres re-innervation onto IHCs because re-innervation occurs when disrupting molecular components in MET apparatus (Corns et al., 2018; O’Connor et al., 2024). However, re-innervation does not correlate to the degree of MET current degeneration with age in C57BL/6N and C57BL/6N with corrected *Cdh23^ahl^* mutation (6N-Repaired, Mianné et al., 2016) (Jeng, Carlton, et al., 2021b). If other aging processes cause efferent re-innervation, CBA/CaJ IHCs should develop efferent contact disregard of their near-normal MET current size.

Efferent terminals were visualized using antibodies against choline acetyltransferase (ChAT) for efferent neurons, and SK2, a small conductance Ca^2+^-activated K^+^ channel that is present in the post-synaptic IHCs. During hair cell maturation, the input from cholinergic efferent fibres activate nicotinic acetylcholine receptors (nAChRs) in hair cells. This triggers the opening of s (SK2) that drive cell hyperpolarisation (Glowatzki & Fuchs, 2000). After hearing onset, mouse IHCs lose contact with efferent neurons and their ability to respond to acetylcholine (Glowatzki & Fuchs, 2000; Marcotti et al., 2004a; Figure 6A). With age, IHCs from C57BL/6 mice establish functional connections with inhibitory efferent neurons (Lauer et al., 2012; Zachary & Fuchs, 2015; Jeng, Carlton, et al., 2021b). In this study, results show that CBA/CaJ mice expressed more SK2 channels with age which juxtapose ChAT positive neuronal terminals (Figure 6B). The number of IHCs with SK2 puncta also significantly increased with age (1 month vs. 21–23 months: P < 0.001, Figure 6C). However, this number was less compared to that from C3H/HeJ mice (CBA/CaJ vs. C3H: P < 0.001; vs. C57BL/6N: P = 0.028, vs. 6N-Repaired: P = 0.076, Figure 6C). CBA/CaJ mice also show significantly less SK2 puncta per cell than C3H/HeJ, indicating that the difference between strains is particularly evident among CBA/CaJ and C3H/HeJ mice (P < 0.001, Figure 6D). At the old ages, CBA/CaJ IHCs are innervated by efferent fibres, suggesting the rewiring process does not depend on the degree of hearing loss nor defects in MET apparatus.

**Figure 6.**
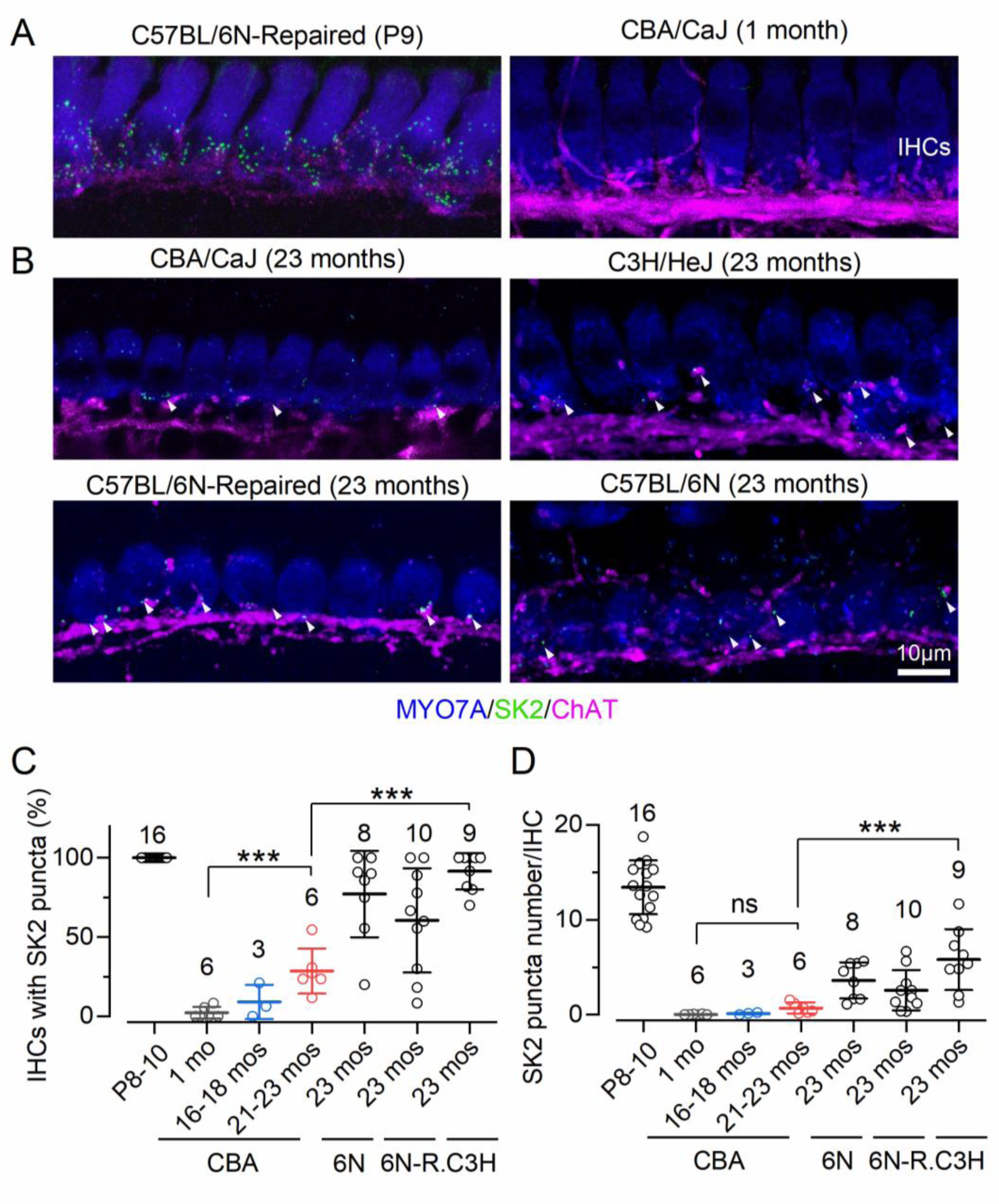
Efferent synapses contacted ageing inner hair cells. Cochlea from a 100 µm region of the apical coil from ageing CBA/CaJ (CBA), C3H/HeJ (C3H), C57BL/6N-Repaired (6N-R) and C57BL/6N (6N) mice were labelled with antibodies against a hair cell marker MYO7A (blue), SK2 channels (green), and ChAT (magenta). *A*, maximum intensity projections of confocal z-stacks from apical-coil IHCs of mice at P9 and 1 month. *B*, maximum intensity projections of confocal z-stacks from apical-coil IHCs of mice at 23 months. *C*, percentage of IHCs showing SK2 puncta with age. *D*, number of SK2 puncta per IHC with age. Data are plotted as mean ± SD. Number of animal is shown above the data. Single counts are plotted as open symbols. *** P < 0.001, one-way ANOVA followed by Tukey’s post-test, ns: not significant.

## Discussion

ARHL is the most prevalent sensory disorder, yet its underlying mechanisms remain elusive, associated with complex genetic and environmental factors. Understanding the onset of ARHL is crucial for developing early treatment interventions that precede the irreversible hearing loss. A primary pathology preceding hearing threshold elevation is synaptopathy, characterized by the age-dependent decline of hair cell ribbon synapses, though the underlying mechanism is unclear. In particular, the relationship between ribbon synapse loss and declining hair cell function requires further investigation. In this study, we aimed to determine whether hair cells exhibit functional degeneration with age. We provide evidence that hair cells become smaller and display a decreased K^+^ current profile at old ages, a functional change that occurs prior to the elevation of hearing thresholds.

We found that the auditory system’s aging process differs significantly between mouse strains. In aging CBA/CaJ mice, hair cells become smaller and express reduced *I*_K_ and *I*_K,f_ (Figure 2), a feature also observed in C57BL but absent in C3H/HeJ mice (Jeng, Johnson, et al., 2020; Jeng, Carlton, et al., 2021b). Consistent with these functional deficits, previous literatures indicates that CBA/CaJ but not C3H mice begin losing synaptic ribbons within similar age period at the same apical cochlear regions (Parthasarathy & Kujawa, 2018; Jeng, Ceriani, et al., 2020; Dörje et al., 2024). This divergence is intriguing because, despite originating from the same initial cross, CBA/CaJ mice exhibit physiological age-related hair cell changes distinct from those in C3H/HeJ mice. These differences underscore the significant influence of genetic background on auditory aging.

Despite changes in the basolateral current profiles, CBA/CaJ did not show a decrease in MET current up to about 20 months of age (Figure 4). This finding is expected given the absence of profound hearing loss at this stage. However, as the total basolateral current responses were smaller with age (Figure 2), the current balance between MET current and basolateral current may not be maintained for a normal resting membrane potential. This is supported by the observation of a more depolarized resting membrane potential in older (16–18 months) compared to young (1 month) IHCs (Figure 5G). This might reflect a change in the operational range and the spontaneous synaptic vesicle release rate at rest for IHCs in aging mice (Tobón & Moser, 2024). Considering *in vivo* hair cells rest at a higher membrane potential and have a larger MET current response due to a lower Ca^2+^ concentration, changes observed *ex vivo* might have a greater physiological impact *in vivo* (Johnson, 2015). Nevertheless, the potential contribution of this depolarization to excitotoxicity remains unknown.

Our results reveal that aging hair cell defects involve changes in ion channels, specifically reduced total basolateral currents and BK currents. Although the reduced cell size correlates with the total current response, it is insufficient to account for the significant reduction in BK channel currents (Figure 2L). We found that the volume of anti-BK channel staining decreased with age, although its expression remained restricted to the neck region of IHCs (Figure 3). This suggests that the protein network at the neck region may be compartmentalized and not simply affected by general cell shrinkage with age. Functionally, since BK currents are crucial for hair cell repolarization, the loss of this current may impair the cell’s activation range and the temporal accuracy of sound transduction (Kurt et al., 2012). This could thus contribute to hidden hearing loss before an obvious elevation of hearing thresholds in aging CBA/CaJ mice. Furthermore, the role of BK channels may extend beyond modulating hearing sensitivity. Mutations in the BK channel gene (*Kcma1*) do not cause hearing threshold changes at young ages but increase susceptibility to noise-induced and age-related hearing loss, suggesting a long-term consequence of BK dysfunction (Rüttiger et al., 2004; Pyott et al., 2007). As BK is also expressed in OHCs at basal regions of the cochlea (Wersinger et al., 2010; Rohmann et al., 2015), they may also be involved in initiating ARHL at higher frequencies. Therefore, our findings support the recent hypothesis that ion channels, such as BK channel, could be a promising therapeutic target for ARHL (Peixoto Pinheiro et al., 2021).

In this study, we found that IHCs in CBA/CaJ mice express SK2 channels localized near efferent fibers. While this organization was previously identified, likely near lateral olivocochlear (LOC) fibers, in other aging mouse models (Lauer et al., 2012; Zachary & Fuchs, 2015; Jeng, Carlton, et al., 2021b), the full impact of re-innervation is still unclear. One hypothesis is that efferent neurons respond to physiological changes in IHC. Nevertheless, it is unlikely that transduction via hair bundles orchestrates this re-innervation, as the MET current did not change with age in CBA/CaJ and 6N-Repaired mice (Jeng, Carlton, et al., 2021b). If re-innervation in aging hair cells is functionally similar to immature hair cells, the efferent modulation of IHCs might influence the organisation of synapses as reported in developmental stages (Johnson et al., 2013; Clause et al., 2017).

Overall, aging hair cells in CBA/CaJ mice exhibit features that recapitulate immature cells, including smaller cell size, reduced expression of BK channels, and re-innervation by efferent neurons. In conclusion, our findings demonstrate that the hair cells of aging CBA/CaJ mice undergo a progressive functional degeneration that precedes the onset of measurable hearing loss, suggesting cellular deficits as a key step in auditory aging pathology.

## Materials and Methods

### Tissue preparation for ex vivo recordings

Patch clamp recordings were performed from hair cells within the 9-12 kHz region of the cochlear apical coil75(Müller et al., 2005). Cochleae were dissected out in extracellular solution composed of (in mM): 135 NaCl, 5.8 KCl, 1.3 CaCl_2_, 0.9 MgCl_2_, 0.7 NaH_2_PO_4_, 5.6 D-glucose, 10 HEPES-NaOH. Sodium pyruvate (2 mM), amino acids and vitamins were added from concentrates (Thermo Fisher Scientific, UK). The pH was adjusted to 7.48 with 1M NaOH, and the final osmolality of the solution was measured (∼306 mmol/kg). The dissected apical coil was then transferred to a microscope chamber and immobilised via a nylon mesh attached to a stainless-steel ring. The chamber (volume 2 ml) was perfused from a peristaltic pump (Cole-Palmer, UK) and mounted on the stage of an upright microscope (Olympus BX51, Japan) with Nomarski Differential Interference Contrast (DIC) optics. The individual stereocilia rows of the hair cells was identified using a 60x water immersion objective and a 15x eyepiece.

### Whole-cell electrophysiology

Patch clamp recordings were performed at room temperature (20-24°C) using an Optopatch amplifier (Cairn Research Ltd, UK). Patch pipettes were pulled from soda glass capillaries and had a typical resistance in extracellular solution of 2-3 MΩ. The patch pipette intracellular solution contained (in mM): 131 KCl, 3 MgCl_2_, 1 EGTA-KOH, 5 Na_2_ATP, 5 HEPES-KOH, 10 Na-phosphocreatine (pH was adjusted with 1M KOH to 7.28, ∼294 mmol/kg). Data acquisition was controlled by pClamp software using Digidata 1440A and 1550 (Molecular Devices, USA). To reduce the electrode capacitance, patch electrodes were coated with surf wax (Mr. Zoggs SexWax, USA). Recordings were low-pass filtered at 2.5 kHz (8-pole Bessel), sampled at 5 kHz and stored on a computer for off-line analysis (Clampfit, Molecular Devices and Origin 2020, OriginLab). For voltage-clamp experiments, membrane potentials were corrected off-line for the residual series resistance after compensation (60–80%) and the liquid junction potential (LJP) of -4 mV, which was measured between electrode and bath solutions. Holding currents were plotted as zero current to allow a better comparison between recordings. Voltage clamp protocols are referred to a holding potential of -84 mV or -64 mV depending on the protocol used.

For mechanoelectrical transducer (MET) current recordings, the hair bundles of hair cells were displaced using a fluid jet from a pipette driven by a 25 mm diameter piezoelectric disc (Corns & Marcotti, 2016; J.-Y. Jeng, Harasztosi, et al., 2021). The pipette was pulled from borosilicate glass to a final overall length of 5.3-5.5 cm. The fluid jet pipette tip had a diameter of 8-10 µm and was positioned near the hair bundles to elicit a maximal MET current. Mechanical stimuli were applied as 50 Hz sinusoids (filtered at 1 kHz, 8-pole Bessel). Prior to the positioning of the fluid jet by the hair bundles, any steady-state pressure was removed by monitoring the movement of debris in front of the pipette. The use of the fluid-jet allows for the efficient displacement of the hair bundles in both the excitatory and inhibitory directions, which is essential to perform reliable measurements of the resting open probability of the MET channels.

### Auditory brainstem responses

Following the onset of anaesthesia (see Ethics statement above) and the loss of the retraction reflex with a toe pinch, mice were placed in a soundproof chamber (MAC-3 acoustic chamber, IAC Acoustic, UK). Male and female mice were placed on a heated mat (37°C) with the animal’s pinna being positioned at a distance of 10 cm from the loudspeaker. Two subdermal electrodes were placed under the skin behind the pinna of each ear (reference and ground electrode), and one electrode half-way between the two pinna on the vertex of the cranium (active electrode) as previously described (Ingham et al., 2011). Sound stimuli were delivered to the mouse ear by a loudspeaker (MF1-S, Multi Field Speaker, Tucker-Davis Technologies, USA), which was calibrated with a low-noise microphone probe system (ER10B+, Etymotic, USA). Experiments were performed using customized software (Ingham et al., 2011) driving an RZ6 auditory processor (Tucker-Davis Technologies). ABR thresholds, which were delivered as white noise clicks and pure tone stimuli of frequencies at 6, 12, 18, 24, 30, 36 and 42 kHz, were defined as the lowest sound level where any recognisable feature of the waveform was visible. Stimulus sound pressure levels were up to 95 dB SPL, were presented in steps of 5 dB SPL (averaged over 256 repetitions). Tone bursts were 5 ms in duration with a 1 ms on/off ramp time presented at a rate of 42.6/sec.

### Scanning electron microscopy (SEM)

The dissected mouse cochleae were initially fixed by a very gentle intra-labyrinthine perfusion of fixative using a pipette tip through the round window. Following perfusion, the cochleae were immersed in the fixative for 2 hrs at room temperature. The fixative contained 2.5% glutaraldehyde in 0.1 M sodium cacodylate buffer and 2 mM CaCl_2_ (pH 7.4). After the fixation, the organ of Corti was exposed by removing the bone from the apical coil of the cochleae, then post-fix in 1% osmium tetroxide in the cacodylate buffer for 1 h. For osmium impregnation, cochleae were incubated in solutions of saturated aqueous thiocarbohydrazide (20 mins) alternating with 1% osmium tetroxide in cacodylate buffer (2 hrs) twice (OTOTO technique). The cochleae were then dehydrated through an ethanol series (70–100%) and critical point dried using CO_2_ as the transitional fluid (Leica EM CPD300), and mounted on specimen stubs using conductive silver paint (Agar Scientific, Stansted, UK). The apical coil of the organ of Corti was examined at 10 kV using a Tescan Vega3 LMU scanning electron microscope at the School of Biosciences Electron Microscopy Facility at the University of Sheffield.

### Immunofluorescence microscopy

The majority of this work was performed by fixing the inner ear with 4% paraformaldehyde in phosphate-buffered saline (PBS, pH 7.4) for 20 minutes at room temperature. Cochleae were washed three times in PBS for 10 minutes and fine dissected for the Organ of Corti. Following dissection, samples were incubated in PBS supplemented with 5% normal goat or horse serum and 0.5% Triton X-100 for 1 hour at room temperature. The samples were immunolabelled with primary antibodies overnight at 37°C, washed three times with PBS and incubated with the secondary antibodies for 1 hour at 37°C. Antibodies were prepared in 1% serum and 0.5% Triton X-100 in PBS. Primary antibodies were: mouse anti-myosin 7a (1:1000, Developmental Studies Hybridoma Bank, #138-1C), rabbit anti-myosin 7a (1:200, Proteus Biosciences, #25-6790), rabbit anti-SK2 (1:500, Sigma-Aldrich, P0483) goat anti-choline acetyltransferase (ChAT, 1:500, Millipore, AB144P), and mouse-IgG1 anti-BK channel (1:200, NeuroMab, 75-408). Secondary antibodies were species appropriate Alexa Fluor secondary antibodies. Samples were mounted in VECTASHIELD (H-1000). The images from the apical cochlear region (8-12 kHz) were captured with a Zeiss LSM 880 Airyscan equipped with Plan-Apochromat 63x Oil DIC M27 objective. The microscope is part of the Wolfson Light Microscope Facility at the University of Sheffield. Image stacks were processed with Fiji ImageJ software (https://imagej.net/Fiji).

### Statistical analysis

Statistical comparisons of means were made by Student’s two-tailed t test or for multiple comparisons, analysis of variance (one-way and two-way ANOVA followed by Tukey’s or Sidak’s post hoc test). Only mean values with a similar variance between groups were compared and values are quoted in text and figures as means ± standard deviation. In the presence of an upper threshold limit of equipment, such as when determining hearing thresholds, non-parametric Kruskal-Wallis ANOVA was used and values are quoted in text and figures as median ± median absolute deviation. P < 0.05 was selected as the criterion for statistical significance.

## Author contribution

All authors helped with the analysis of the data. J-YJ conceived and coordinated the study. All authors approved the final version of the manuscript. All authors agree to be accountable for all aspects of the work in ensuring that questions related to the accuracy or integrity of any part of the work are appropriately investigated and resolved.

## Acknowledgments

The authors would like to acknowledge the Wolfson Light Microscopy Facility and the Biosciences Electron Microscopy Facility at the University of Sheffield for their guidance and services.

## Ethics Statement

The animal work was licensed by the UK Home Office under the Animals (Scientific Procedures) Act 1986 (PPL_PCC8E5E93) and was approved by the University of Sheffield Ethical Review Committee (180626_Mar). Both males and females where used for this study. For ex vivo experiments mice were killed by cervical dislocation followed by decapitation. For in vivo auditory brainstem responses (ABRs) and distortion product otoacoustic emissions (DPOAEs) mice were anesthetized using intraperitoneal injection of ketamine (100 mg/Kg body weight, Fort Dodge Animal Health, USA) and xylazine (10 mg/Kg, Rompun, Bayer HealthCare LLC, USA). At the end of the ABR recordings, mice were either culled by cervical dislocation or recovered from anesthesia with intraperitoneal injection of atipamezole (1 mg/Kg, Antisedan, Orion Corporation, Finland).

## References

Assad, J. A., Shepherd, G. M., & Corey, D. P. (1991). Tip-link integrity and mechanical transduction in vertebrate hair cells. Neuron, 7(6), 985–994.

Beurg, M., Cui, R., Goldring, A. C., Ebrahim, S., Fettiplace, R., & Kachar, B. (2018). Variable number of TMC1-dependent mechanotransducer channels underlie tonotopic conductance gradients in the cochlea. Nature Communications, 9, 2185. 10.1038/s41467-018-04589-8

Beurg, M., Fettiplace, R., Nam, J.-H., & Ricci, A. J. (2009). Localization of inner hair cell mechanotransducer channels using high-speed calcium imaging. Nature Neuroscience, 12(5), 553. 10.1038/nn.2295

Cieza, A., Causey, K., Kamenov, K., Hanson, S. W., Chatterji, S., & Vos, T. (2021). Global estimates of the need for rehabilitation based on the Global Burden of Disease study 2019: A systematic analysis for the Global Burden of Disease Study 2019. *Lancet (London*, England), 396(10267), 2006–2017. 10.1016/S0140-6736(20)32340-0

Clause, A., Lauer, A. M., & Kandler, K. (2017). Mice Lacking the Alpha9 Subunit of the Nicotinic Acetylcholine Receptor Exhibit Deficits in Frequency Difference Limens and Sound Localization. Frontiers in Cellular Neuroscience, 11, 167. 10.3389/fncel.2017.00167

Corns, L. F., Johnson, S. L., Roberts, T., Ranatunga, K. M., Hendry, A., Ceriani, F., Safieddine, S., Steel, K. P., Forge, A., Petit, C., Furness, D. N., Kros, C. J., & Marcotti, W. (2018). Mechanotransduction is required for establishing and maintaining mature inner hair cells and regulating efferent innervation. Nature Communications, 9(1), 4015. 10.1038/s41467-018-06307-w

Corns, L. F., & Marcotti, W. (2016). Piezo1 haploinsufficiency does not alter mechanotransduction in mouse cochlear outer hair cells. Physiological Reports, 4(3), e12701. 10.14814/phy2.12701

Dörje, N. M., Shvachiy, L., Kück, F., Outeiro, T. F., Strenzke, N., Beutner, D., & Setz, C. (2024). Age-related alterations in efferent medial olivocochlear-outer hair cell and primary auditory ribbon synapses in CBA/J mice. Frontiers in Cellular Neuroscience, 18. 10.3389/fncel.2024.1412450

Gacek, R. R., & Schuknecht, H. F. (1969). Pathology of Presbycusis. International Audiology, 8(2– 3), 199–209. 10.3109/05384916909079061

Gioacchini, F. M., Pisani, D., Viola, P., Astorina, A., Scarpa, A., Libonati, F. A., Tulli, M., Re, M., & Chiarella, G. (2023). Diabetes Mellitus and Hearing Loss: A Complex Relationship. Medicina, 59(2), 269. 10.3390/medicina59020269

Glowatzki, E., & Fuchs, P. A. (2000). Cholinergic Synaptic Inhibition of Inner Hair Cells in the Neonatal Mammalian Cochlea. Science, 288(5475), 2366–2368. 10.1126/science.288.5475.2366

Gorur-Shandilya, S., Marder, E., & O’Leary, T. (2020). Activity-dependent compensation of cell size is vulnerable to targeted deletion of ion channels. Scientific Reports, 10(1), 15989. 10.1038/s41598-020-72977-6

Ingham, N. J., Pearson, S., & Steel, K. P. (2011). Using the Auditory Brainstem Response (ABR) to Determine Sensitivity of Hearing in Mutant Mice. Current Protocols in Mouse Biology, 1(2), 279–287. 10.1002/9780470942390.mo110059

Jeng, J., Carlton, A., Johnson, S. L., Brown, S. D. M., Holley, M. C., Bowl, M. R., & Marcotti, W. (2020). Biophysical and morphological changes in inner hair cells and their efferent innervation in the ageing mouse cochlea. The Journal of Physiology, JP280256. 10.1113/JP280256

Jeng, J.-Y., Carlton, A. J., Goodyear, R. J., Chinowsky, C., Ceriani, F., Johnson, S. L., Sung, T.-C., Dayn, Y., Richardson, G. P., Bowl, M. R., Brown, S. D. M., Manor, U., & Marcotti, W. (2022). AAV-mediated rescue of Eps8 expression in vivo restores hair-cell function in a mouse model of recessive deafness. Molecular Therapy - Methods & Clinical Development, 26, 355–370. 10.1016/j.omtm.2022.07.012

Jeng, J.-Y., Carlton, A. J., Johnson, S. L., Brown, S. D. M., Holley, M. C., Bowl, M. R., & Marcotti, W. (2021). Biophysical and morphological changes in inner hair cells and their efferent innervation in the ageing mouse cochlea. The Journal of Physiology, 599(1), 269–287. 10.1113/JP280256

Jeng, J.-Y., Ceriani, F., Olt, J., Brown, S. D. M., Holley, M. C., Bowl, M. R., Johnson, S. L., & Marcotti, W. (2020). Pathophysiological changes in inner hair cell ribbon synapses in the ageing mammalian cochlea. The Journal of Physiology, 598(19), 4339–4355. 10.1113/JP280018

Jeng, J.-Y., Harasztosi, C., Carlton, A. J., Corns, L. F., Marchetta, P., Johnson, S. L., Goodyear, R. J., Legan, K. P., Rüttiger, L., Richardson, G. P., & Marcotti, W. (2021). MET currents and otoacoustic emissions from mice with a detached tectorial membrane indicate the extracellular matrix regulates Ca2+ near stereocilia. The Journal of Physiology, 599(7), 2015–2036. 10.1113/JP280905

Jeng, J.-Y., Johnson, S. L., Carlton, A. J., Tomasi, L. D., Goodyear, R. J., Faveri, F. D., Furness, D. N., Wells, S., Brown, S. D. M., Holley, M. C., Richardson, G. P., Mustapha, M., Bowl, M. R., & Marcotti, W. (2020). Age-related changes in the biophysical and morphological characteristics of mouse cochlear outer hair cells. The Journal of Physiology, 598(18), 3891– 3910. 10.1113/JP279795

Johnson, S. L. (2015). Membrane properties specialize mammalian inner hair cells for frequency or intensity encoding. eLife, 4, e08177. 10.7554/eLife.08177

Johnson, S. L., Wedemeyer, C., Vetter, D. E., Adachi, R., Holley, M. C., Elgoyhen, A. B., & Marcotti, W. (2013). Cholinergic efferent synaptic transmission regulates the maturation of auditory hair cell ribbon synapses. Open Biology, 3(11), 130163. 10.1098/rsob.130163

Kharkovets, T., Dedek, K., Maier, H., Schweizer, M., Khimich, D., Nouvian, R., Vardanyan, V., Leuwer, R., Moser, T., & Jentsch, T. J. (2006). Mice with altered KCNQ4 K+ channels implicate sensory outer hair cells in human progressive deafness. The EMBO Journal, 25(3), 642–652. 10.1038/sj.emboj.7600951

Kros, C. J., Ruppersberg, J. P., & Rüsch, A. (1998). Expression of a potassium current in inner hair cells during development of hearing in mice. Nature, 394(6690), 281–284. 10.1038/28401

Kurt, S., Sausbier, M., Rüttiger, L., Brandt, N., Moeller, C. K., Kindler, J., Sausbier, U., Zimmermann, U., Straaten, H. van, Neuhuber, W., Engel, J., Knipper, M., Ruth, P., & Schulze, H. (2012). Critical role for cochlear hair cell BK channels for coding the temporal structure and dynamic range of auditory information for central auditory processing. The FASEB Journal, 26(9), 3834–3843. 10.1096/fj.11-200535

Lauer, A. M., Fuchs, P., Ryugo, D. K., & Francis, H. W. (2012). Efferent synapses return to inner hair cells in the aging cochlea. Neurobiology of Aging, 33(12), 2892–2902. 10.1016/j.neurobiolaging.2012.02.007

Liberman, M. C., & Kujawa, S. G. (2017). Cochlear synaptopathy in acquired sensorineural hearing loss: Manifestations and mechanisms. Hearing Research, 349, 138–147. 10.1016/j.heares.2017.01.003

Lingle, C. J., Martinez-Espinosa, P. L., Yang-Hood, A., Boero, L. E., Payne, S., Persic, D., V-Ghaffari, B., Xiao, M., Zhou, Y., Xia, X.-M., Pyott, S. J., & Rutherford, M. A. (2019). LRRC52 regulates BK channel function and localization in mouse cochlear inner hair cells. Proceedings of the National Academy of Sciences of the United States of America, 116(37), 18397–18403. 10.1073/pnas.1907065116

Marcotti, W., Johnson, S. L., & Kros, C. J. (2004a). A transiently expressed SK current sustains and modulates action potential activity in immature mouse inner hair cells. The Journal of Physiology, 560(Pt 3), 691–708. 10.1113/jphysiol.2004.072868

Marcotti, W., Johnson, S. L., & Kros, C. J. (2004b). Effects of intracellular stores and extracellular Ca2+ on Ca2+-activated K+ currents in mature mouse inner hair cells. The Journal of Physiology, 557(Pt 2), 613–633. 10.1113/jphysiol.2003.060137

Marcotti, W., & Kros, C. J. (1999). Developmental expression of the potassium current IK,n contributes to maturation of mouse outer hair cells. The Journal of Physiology, 520(3), 653– 660. 10.1111/j.1469-7793.1999.00653.x

Mianné, J., Chessum, L., Kumar, S., Aguilar, C., Codner, G., Hutchison, M., Parker, A., Mallon, A.- M., Wells, S., Simon, M. M., Teboul, L., Brown, S. D. M., & Bowl, M. R. (2016). Correction of the auditory phenotype in C57BL/6N mice via CRISPR/Cas9-mediated homology directed repair. Genome Medicine, 8, 16. 10.1186/s13073-016-0273-4

O’Connor, A. P., Amariutei, A. E., Zanella, A., Hool, S. A., Carlton, A. J., Kong, F., Saenz-Roldan, M., Jeng, J.-Y., Lecomte, M.-J., Johnson, S. L., Safieddine, S., & Marcotti, W. (2024). In vivo AAV9-Myo7a gene rescue restores hearing and cholinergic efferent innervation in inner hair cells. JCI Insight, 9(23), e182138. 10.1172/jci.insight.182138

Ohlemiller, K. K., Dahl, A. R., & Gagnon, P. M. (2010). Divergent Aging Characteristics in CBA/J and CBA/CaJ Mouse Cochleae. JARO: Journal of the Association for Research in Otolaryngology, 11(4), 605–623. 10.1007/s10162-010-0228-1

Oliver, D., Knipper, M., Derst, C., & Fakler, B. (2003). Resting potential and submembrane calcium concentration of inner hair cells in the isolated mouse cochlea are set by KCNQ-type potassium channels. The Journal of Neuroscience: The Official Journal of the Society for Neuroscience, 23(6), 2141–2149.

Parthasarathy, A., & Kujawa, S. G. (2018). Synaptopathy in the Aging Cochlea: Characterizing Early-Neural Deficits in Auditory Temporal Envelope Processing. The Journal of Neuroscience, 38(32), 7108–7119. 10.1523/JNEUROSCI.3240-17.2018

Peixoto Pinheiro, B., Vona, B., Löwenheim, H., Rüttiger, L., Knipper, M., & Adel, Y. (2021). Age-related hearing loss pertaining to potassium ion channels in the cochlea and auditory pathway. Pflugers Archiv, 473(5), 823–840. 10.1007/s00424-020-02496-w

Pyott, S. J., Meredith, A. L., Fodor, A. A., Vázquez, A. E., Yamoah, E. N., & Aldrich, R. W. (2007). Cochlear Function in Mice Lacking the BK Channel α, β1, or β4 Subunits. Journal of Biological Chemistry, 282(5), 3312–3324. 10.1074/jbc.M608726200

Rohmann, K. N., Wersinger, E., Braude, J. P., Pyott, S. J., & Fuchs, P. A. (2015). Activation of BK and SK Channels by Efferent Synapses on Outer Hair Cells in High-Frequency Regions of the Rodent Cochlea. The Journal of Neuroscience, 35(5), 1821–1830. 10.1523/JNEUROSCI.2790-14.2015

Rüttiger, L., Sausbier, M., Zimmermann, U., Winter, H., Braig, C., Engel, J., Knirsch, M., Arntz, C., Langer, P., Hirt, B., Müller, M., Köpschall, I., Pfister, M., Münkner, S., Rohbock, K., Pfaff, I., Rüsch, A., Ruth, P., & Knipper, M. (2004). Deletion of the Ca2+-activated potassium (BK) alpha-subunit but not the BKbeta1-subunit leads to progressive hearing loss. Proceedings of the National Academy of Sciences of the United States of America, 101(35), 12922–12927. 10.1073/pnas.0402660101

Sergeyenko, Y., Lall, K., Liberman, M. C., & Kujawa, S. G. (2013). Age-Related Cochlear Synaptopathy: An Early-Onset Contributor to Auditory Functional Decline. The Journal of Neuroscience, 33(34), 13686–13694. 10.1523/JNEUROSCI.1783-13.2013

Sha, S.-H., Kanicki, A., Dootz, G., Talaska, A. E., Halsey, K., Dolan, D., Altschuler, R., & Schacht, J. (2008). Age-related auditory pathology in the CBA/J mouse. Hearing Research, 243(1), 87–94. 10.1016/j.heares.2008.06.001

Solsona, C., Innocenti, B., & Fernández, J. M. (1998). Regulation of Exocytotic Fusion by Cell Inflation. Biophysical Journal, 74(2), 1061–1073. 10.1016/S0006-3495(98)74030-5

Spongr, V. P., Flood, D. G., Frisina, R. D., & Salvi, R. J. (1997). Quantitative measures of hair cell loss in CBA and C57BL/6 mice throughout their life spans. The Journal of the Acoustical Society of America, 101(6), 3546–3553.

Tang, D., Tran, Y., Dawes, P., & Gopinath, B. (2023). A Narrative Review of Lifestyle Risk Factors and the Role of Oxidative Stress in Age-Related Hearing Loss. Antioxidants, 12(4), 878. 10.3390/antiox12040878

Tobón, L. M. J., & Moser, T. (2024). Bridging the gap between presynaptic hair cell function and neural sound encoding. eLife, 12. 10.7554/eLife.93749.3

Wersinger, E., McLean, W. J., Fuchs, P. A., & Pyott, S. J. (2010). BK Channels Mediate Cholinergic Inhibition of High Frequency Cochlear Hair Cells. PLoS ONE, 5(11). 10.1371/journal.pone.0013836

World report on hearing. (2021). World Health Organization.

Wu, P. Z., Liberman, L. D., Bennett, K., de Gruttola, V., O’Malley, J. T., & Liberman, M. C. (2018). Primary Neural Degeneration in the Human Cochlea: Evidence for Hidden Hearing Loss in the Aging Ear. Neuroscience. 10.1016/j.neuroscience.2018.07.053

Wu, P.-Z., O’Malley, J. T., Gruttola, V. de, & Liberman, M. C. (2021). Primary Neural Degeneration in Noise-Exposed Human Cochleas: Correlations with Outer Hair Cell Loss and Word-Discrimination Scores. Journal of Neuroscience, 41(20), 4439–4447. 10.1523/JNEUROSCI.3238-20.2021

Yamasoba, T., Lin, F. R., Someya, S., Kashio, A., Sakamoto, T., & Kondo, K. (2013). Current concepts in age-related hearing loss: Epidemiology and mechanistic pathways. Hearing Research, 303, 30–38. 10.1016/j.heares.2013.01.021

Zachary, S. P., & Fuchs, P. A. (2015). Re-Emergent Inhibition of Cochlear Inner Hair Cells in a Mouse Model of Hearing Loss. The Journal of Neuroscience, 35(26), 9701–9706. 10.1523/JNEUROSCI.0879-15.2015

